# Morphine-induced side effects can be differentially modulated by cannabidiol in male and female rats

**DOI:** 10.1101/2024.01.22.576721

**Authors:** Carlos Henrique Alves Jesus, Jaqueline Volpe, Bruna Bittencourt Sotomaior, Maria Augusta Ruy Barbosa, Matheus Vinicius Ferreira, Fernanda Fiatcoski, Karina Genaro, José Alexandre de Souza Crippa, Dênio Emanuel Pires Souto, Joice Maria da Cunha

## Abstract

Opioid use disorder (OUD) is a public health problem that includes symptoms such as withdrawal syndrome and opioid-induced hyperalgesia (OIH). Currently, drugs to treat side effects of opioids also have undesirable effects, which lead to limitations. This study investigated the effect of a treatment with cannabidiol (CBD) in morphine-induced hyperalgesia and withdrawal signs in morphine-dependent rats. Male and female rats were submitted to morphine-induced physical dependence protocol consisting of a twice daily treatment with morphine (7.89 mg/kg, 1ml/kg, s.c.) for 10 days. Nociception was measured using the hot plate test and morphine-induced thermal hyperalgesia was equally achieved following 7-10 days of morphine administration in male and female rats. Repeated treatment with CBD (30 mg/kg) was sufficient to prevent thermal hyperalgesia in male and female rats. Subsequently, rats received an acute administration of naloxone (2 mg/kg. s.c.), 90 minutes after the morphine treatment on day 11, the number of withdrawal signs was scored. Rats that received treatment exclusively with morphine presented significant withdrawal signs compared to control (Water). Morphine-dependent female rats showed a prevalent stereotyped behavior of rearing, whereas male rats had the sign of teeth chattering as the most preeminent. Treatment with CBD on day 11 partially attenuated the withdrawal signs in morphine-dependent male rats, but not female rats. Altogether, our data provide evidence of an anti-hyperalgesic effect of CBD in rats. Male and female rats treated chronically with morphine exhibited withdrawal signs in different ratios, indicating sex-differences in withdrawal behavior and CBD attenuated withdrawal signs in a sex-dependent manner.

## 1. Introduction

Chronic use of opioids is associated with several side effects, including tolerance, hyperalgesia, dependence, and abuse [1]. These problematic consequences are defined as opioid use disorder (OUD), a major public health concern that is estimated to affect approximately 26.8 million people worldwide [2] and annually, an increasing number of deaths is attributable to opioid use [3,4]. Clinical and non-clinical studies have shown that females are more sensitive to the rewarding effects of addictive drugs [5–7] and present higher severity of withdrawal symptoms [8,9], compared to males. Importantly, females were reported to transition faster than males to OUD [10]. It has also been reported that females show a higher severity of symptoms at late morphine withdrawal. The severity of symptoms was positively correlated with the phosphorylated CREB in the ventral tegmental area of the brain, a key area for the reward system[9]. Chronic treatment with morphine has also been associated with a selective internalization of the μ-opioid receptor in the locus coeralueus of male, but not female rats. In addition, estrogen has show to potentiate a switch μ-opioid receptors from a coupling with Gαi/o to a coupling with Gβs proteins in female rats[11]. Despite the sex-related differences, the preclinical research in OUD is still performed predominantly in males.

OUD leads to severe symptoms including tolerance, withdrawal syndrome, opioid-induced hyperalgesia (OIH) and others [1]. The mechanism behind OIH is complex and involves chemical changes in the central nervous system [12]. Although the prevalence and epidemiology of OIH is not well documented [13], OIH is not a rare complication of opioid use and the results vary substantially in the basic literature [14,15].

Pharmacological strategies have been used to minimize the side effects of opioids. However, currently available treatments also have undesirable effects, which lead to potential limitations to their use [16,17]. In recent years, the growth in public support for cannabis legalization and decriminalization has shown the therapeutic potential for cannabis derivatives in neuropsychiatry disorders, including OUD [18]. The endocannabinoid and the opioid system, including its receptors, have shown to interact, and are commonly distributed in areas of the brain (i.e., periaqueductal gray, locus coeruleus, ventral tegmental area and others). For instance, studies have demonstrated that the modulation of CB_1_ receptors can attenuate the development of morphine-induced conditioned place preference[19], whereas the modulation of μ-opioid receptors blocks the conditioned place prefence induce by tetrahydrocannabinol[20]. However, there is non clinical and clinical data on the role of CB_1_ receptors and its agonists on the management of opioid withdrawal, and it shows that CB_1_ agonism can enhance the rewarding properties of opioids and the severity of withdrawal symptoms[21–23]. Thus, the elucidation of the role of CB_1_ receptors in opioid withdrawal, and the use of ligants that do not bind to CB_1_ receptors is of importance for the management of opioid withdrawal.

Cannabidiol (CBD), the second most prevalent compound present in the *Cannabis* plant, has shown to produce a wide range of therapeutic effects such as anti-inflammatory, antioxidant [24], anxiolytic and antidepressant [25,26] and efficacy in substance use disorder [27]. CBD does not show to interact as a direct agonist or antagonist on cannabinoid receptors, but rather as an allosteric modulator in cannabinoid and opioid receptors [28]. In addition, CBD does not show reinforcing effects by itself, and can also reduce the rewarding characteristic of drugs of abuse, such as cocaine and opioids, by mechanisms involving 5-HT1A and TRPV1 receptors, for instance [29–31].

A recent clinical study has shown that CBD may reduce cue-induced craving and anxiety in abstinent individuals, mostly men, with heroin use disorder [32]. Moreover, non-clinical studies have pointed to potential effects of CBD on opioid addictive behaviors [33–35]. In addition, it is very common for opioid users to prepare their injections from commercial tablets designed for oral administration. However, injecting solutions made from tablets involve high risk of embolism and other complications [36]. Thus, improved preparation approaches, such as cold and lukewarm filtration of morphine tablets, has been applied to reduce harm of injection[37].

However, little is known about the effects of CBD in the morphine-induced hyperalgesia, in the opioid withdrawal signs, and in the sex-dependent effects related to its therapeutic efficacy. To offset part of this shortcoming, we carried out experiments to induce morphine side effects by injecting a solution made by cold extraction of commercial morphine tablets, and we aimed to compare the effect of CBD in male and female rats tested in the hot plate test to investigate morphine-induced hyperalgesia. Subsequently, precipitated opioid withdrawal signs were investigated.

## 2. Materials and Methods

### 2.1. Animals

Male and female Wistar rats (200-250 g) were provided by the Federal University of Parana colony and placed in plastic cages (41cmx32cmx16.5cm). The animals were maintained in standard conditions of environment with appropriate temperature (21± 2 °C) and illumination cycle (12 h light/12 h dark), with food and water ad libitum. All experimental procedures and protocols were previously approved by the Federal University of Paraná Institutional Committee for the Ethical Use of Animals (CEUA/BIO-UFPR; authorization #1415). This study was performed in accordance with the ethical guidelines of Brazilian legislation on animal welfare following the ARRIVE guideline. All efforts were made to minimize animal suffering and the number of animals used.

### 2.2. Drugs

Cannabidiol (CBD; 3, 10 or 30 mg/kg, i.p., volume injection 1 ml/kg) 99,6% pure (without any other cannabinoid) was kindly supplied by BSPG-Pharm, Sandwich, United Kingdom. CBD was freshly diluted in a solution of 1:3:16 of tween 80, ethanol and saline. Naloxone (Sigma Aldrich, St. Louis, Missouri, United States) was freshly diluted in saline. The morphine solution used to induce physical dependence and hyperalgesia was prepared by cold extraction of 30 mg tablets of morphine sulfate (Cristália, Itapira, São Paulo, Brazil), carried by the following methodology: several tablets were first crushed using a porcelain mortar and pestle. The resulting powder was mixed with a quantity of distilled water to initially produce a 20 mg/mL morphine solution. This solution was stirred for 20 minutes assisted by an ultrasonic bath, followed by a filtration with a 0.45 µm Durapore membrane filter (Millipore, São Paulo, Brazil). Finally, the solution volume was adjusted with distilled water to obtain an estimated concentration of 10 mg/mL [36]. The final solution concentration was confirmed by UV-Vis spectroscopy (UV-2401 PC, Shimadzu), since morphine has a characteristic absorption band at 285 nm, by using a standard addition method [38]. The standard addition method consisted of adding a predetermined standard morphine solution to the extracted solution diluted by 500x. The standard used was morphine sulfate (Merck S.A, São Paulo, Brazil). Figure 1 displays morphine calibration curve (n=9) at 285 nm absorbance (panel A), and the UV-Vis spectra of one of its replicates, with concentration ranging from 0.01 to 0.1 mg/mL (panel B). The method, previously tested by McLean et al.[36], allowed to obtain similar final concentrations of morphine (7.89 ± 0.06 mg/mL) from distinct extractions.

**Figure 1.**
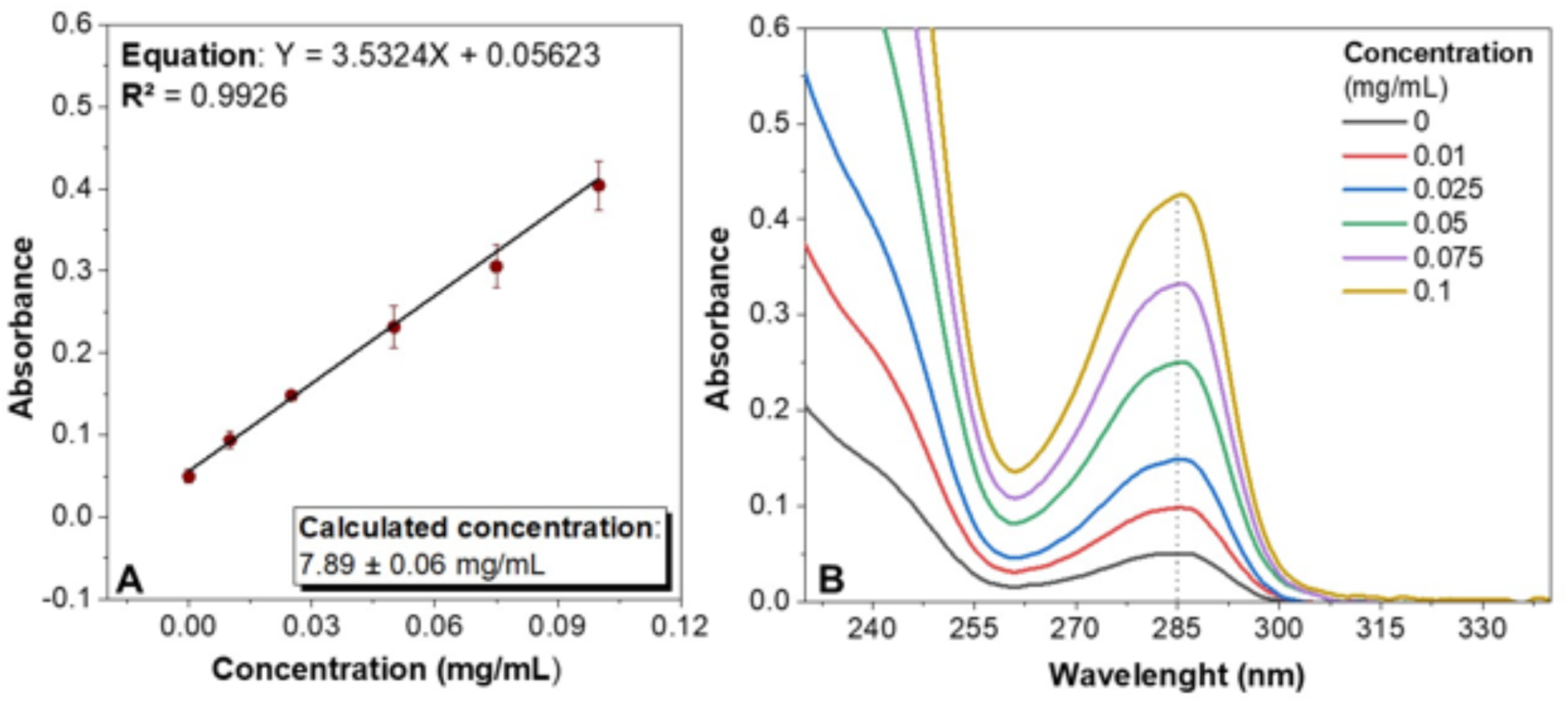
Morphine calibration curve (n=9) at 285 nm absorbance (A), and the UV-Vis spectra of one of its replicates, with concentration ranging from 0.01 to 0.1 mg/mL (B).

### 2.3. Morphine-induced physical dependence and hyperalgesia protocol

Morphine-induced dependence and hyperalgesia were induced as previously described [39,40]. Briefly, all animals were allocated in the laboratory facilities to get used to the environment for 7 days. During acclimation, animals were submitted to daily handling to get used to the experimenter. Male and female rats were made dependent on morphine sulfate with a routine protocol of two subcutaneous (s.c; volume injection 1 ml/kg) injections of morphine (Mor group) twice daily (8:00 am and 6:00 pm) for 10 days (Figure 2A). Rats in the control group received twice-daily injections of distilled water (water group).

**Figure 2.**
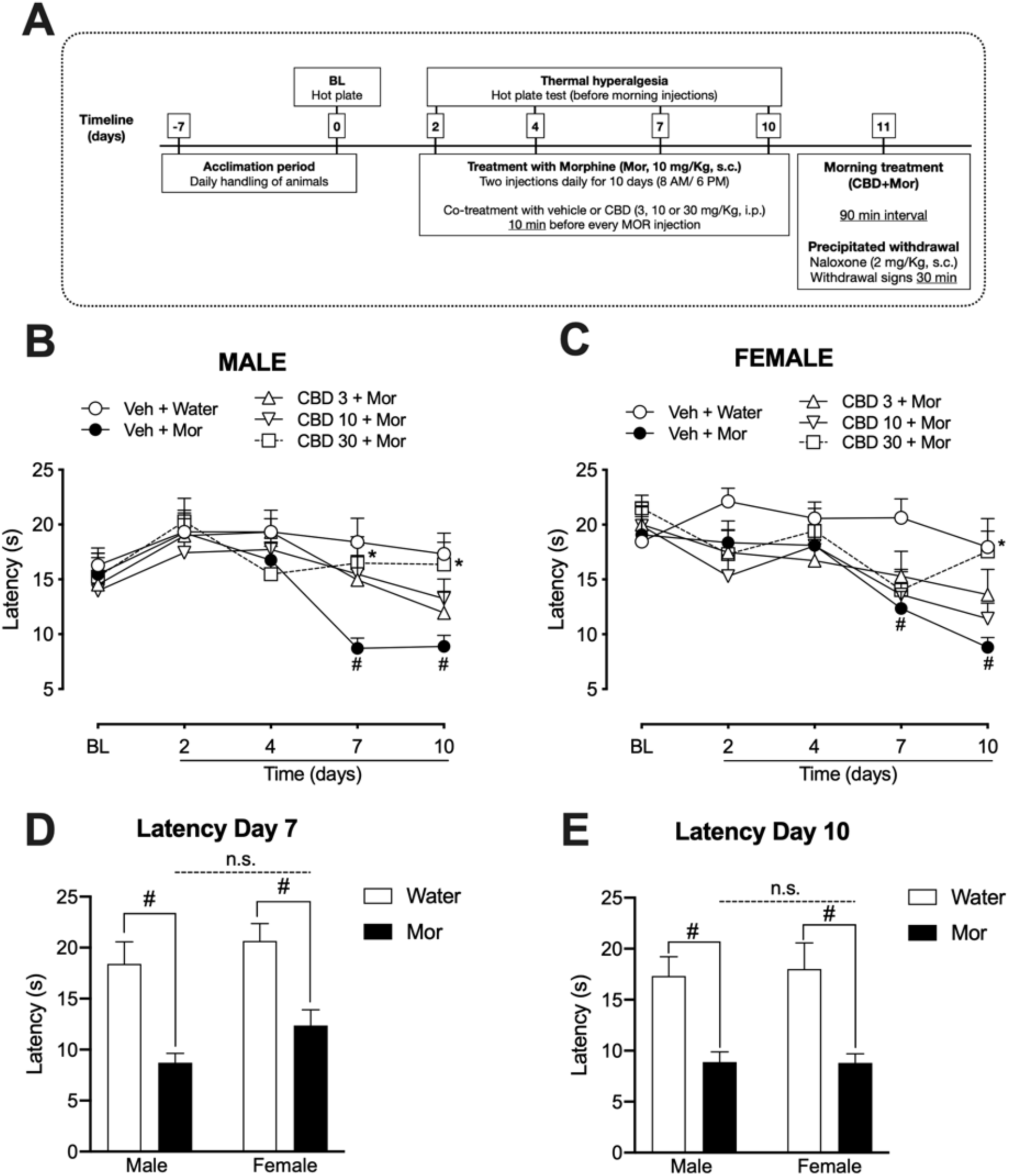
(A) Timeline of the experimental protocols. (B) Effect of CBD treatment in morphine-induced hyperalgesia in male and (B) female rats (C) and evaluation of thermal latency in male and female rats treated with morphine or water on (D) day 7 and (E) day 10 after the beginning of treatments. Groups of male and female Wistar rats treated with CBD (3, 10 or 30 mg/kg; i.p.) before every injection of morphine (s.c.) for 10 days were evaluated in the hot plate test before (baseline, BL), and 2, 4, 7 and 10 days after initial treatment with morphine. Data are expressed as mean ± SEM, n=8-11 per group. # indicates p<0.05 when compared to the respective group treated with water (Veh+Water). * indicates p<0.05 when compared to the respective group treated with morphine (Veh+Mor). Two-way ANOVA followed by Bonferroni’s multiple comparison test.

### 2.4. Hot plate test

According to Pacheco et al. [41], the hot plate test was performed using a hot plate apparatus (Ugo Basile SRL), with the temperature maintained at 50 ± 1°C and a cutoff time of 25 seconds to prevent skin damage. The latency (in seconds) for animals to display behaviors such as licking/flinching of the fore and hind paws or jumping was measured and used to evaluate development of morphine-induced thermal hyperalgesia in male and female rats. Latency was measured before the beginning of treatments with morphine (baseline) and 2, 4, 7 and 10 days after the beginning of treatments (Figure 2A). Treatment with CBD (3, 10 or 30 mg/kg; volume injection 1 ml/kg) was given 10 minutes before each morphine injection, to evaluate the effect of a co-treatment with CBD on the development of morphine-induced hyperalgesia. To avoid the acute antinociceptive effects of treatments (morphine or CBD), the hot plate test was performed before the morning treatments on the specified days. Hot plate tests were performed by an experienced experimenter blind to the treatments.

### 2.5. Naloxone precipitated withdrawal signs

Withdrawal susceptibility was assessed by administration of the selective μ-opioid receptor antagonist naloxone (2 mg/kg; volume injection 1 ml/kg). On the morning of the 11th day, all animals received a single dose of morphine. Fifteen (15) minutes before the evaluation of drug withdrawal, rats were placed in acrylic boxes to acclimate. Ninety minutes after the last morphine treatment, male and female rats received naloxone injection and were immediately placed back in the test box. The number of withdrawal signs such as rearing, teeth chattering, body tremors, defecation (number of fecal boli), digging, sniffing, grooming, and jumping was counted for 30 minutes. Groups of male and female rats were treated with CBD (3, 10 or 30 mg/kg; volume injection 1 ml/kg) 10 minutes before the morphine injection one more time on the 11^th^ day of the protocol, to evaluate the effect of an acute treatment with CBD on naloxone precipitated withdrawal behavior (Figure 2A). The doses of CBD used in the protocol were based on previous studies [26,42,43].

### 2.6. Statistical Analysis

Statistical differences were determined by ANOVAs using GraphPad Prism 8.0 (GraphPad Software, Inc., San Diego, CA, USA). For assessment of behavioral experiments, all data are expressed as mean + standard error. Two-way repeated-measures ANOVA with Bonferroni’s multiple comparisons test was used to assess the effect of treatments, time and interaction between factors. The estimated percentages of each different withdrawal sign calculated from the total number of withdrawal signs during the test period for male and female rats (dependent group only) was compared by unpaired Student’s t-test. One-Way ANOVA followed by the post-hoc analysis of Bonferroni was used to compare the number of withdrawal signs between groups treated with CBD or vehicle. In all cases, the threshold for significance was p < 0.05.

## 3. Results

Figure 2 shows the latency to paw withdrawal evaluated in male (panel B) and female rats (panel C) treated with CBD and morphine. Two-way ANOVA indicated no significant effects of treatment [F(4, 44) = 1.783; p= 0.1494] and interaction between time and treatment [F(16, 176) = 1.441; p= 0.1272], but significant effect of time [F(4, 176) = 8.729; p< 0.0001] in male rats. In female rats, two-way ANOVA indicated no significant effects of treatment [F(4, 40) = 2.562; p= 0.0531] and interaction between time and treatment [F(16, 160) = 1.386; p= 0.1545], but significant effect of time [F(4, 160) = 9.662; p< 0.0001].

Bonferroni’s multiple comparison test showed that male rats treated with morphine (Veh+Mor) had a reduced latency to paw withdrawal in the hot plate test, when compared to its respective water treated group (Veh+Water) on days 7 (p=0.0019) and 10 after the initial treatment (p=0.0088), suggesting the development of OIH. Repeated treatment with CBD at the dose of 30 mg/kg (CBD 30 + Mor) reduced OIH in male rats on day 7 (p=0.0242) and 10 (p=0.0322; Figure 2 panel B).

Bonferroni’s multiple comparison test also revealed that female rats treated with morphine (Veh+Mor) showed reduced latency to paw withdrawal in the hot plate test, when compared to its respective water treated group (Veh+Water) on days 7 (p=0.0202) and 10 after the initial treatment (p=0.0078), which also suggests the development of OIH. Repeated treatment with CBD (30 mg/kg, CBD 30 + Mor) reduced OIH in female rats on day 10 (p=0.0057; Figure 2 panel C).

Figure 2 panel D shows the latency to paw withdrawal evaluated on day 7 after the beginning of treatments, only in the control groups of male and female rats treated with vehicle or morphine and not treated with CBD. Two-way ANOVA indicated significant effect of treatment [F(1, 32) = 28.76; p<0.0001], but not sex [F(1, 32) = 3.10; p=0.0878] or an interaction between these factors [F(1, 32) = 0.16; p= 0.1694]. Bonferroni’s multiple comparison test indicated no significant differences between female and male groups treated with morphine (Mor) 7 days after the beginning of treatments. As for 10 days after the beginning of treatments (Figure 2, panel E), two-way ANOVA also indicated only an effect of treatment [F(1, 32) = 26.64; p<0.0001], suggesting that thermal hyperalgesia was developed at the same time frame in male and female rats treated twice daily with morphine.

As shown in Table 1 and Figure 3, morphine-dependent male and female rats showed significant signs of withdrawal after injection of naloxone (2 mg/kg, s.c.) when compared to their respective control groups treated with water. Two-way ANOVA revealed significant differences in treatment and/or sex and/or interaction between these factors in the different signs of withdrawal evaluated (Table 1).

**Figure 3.**
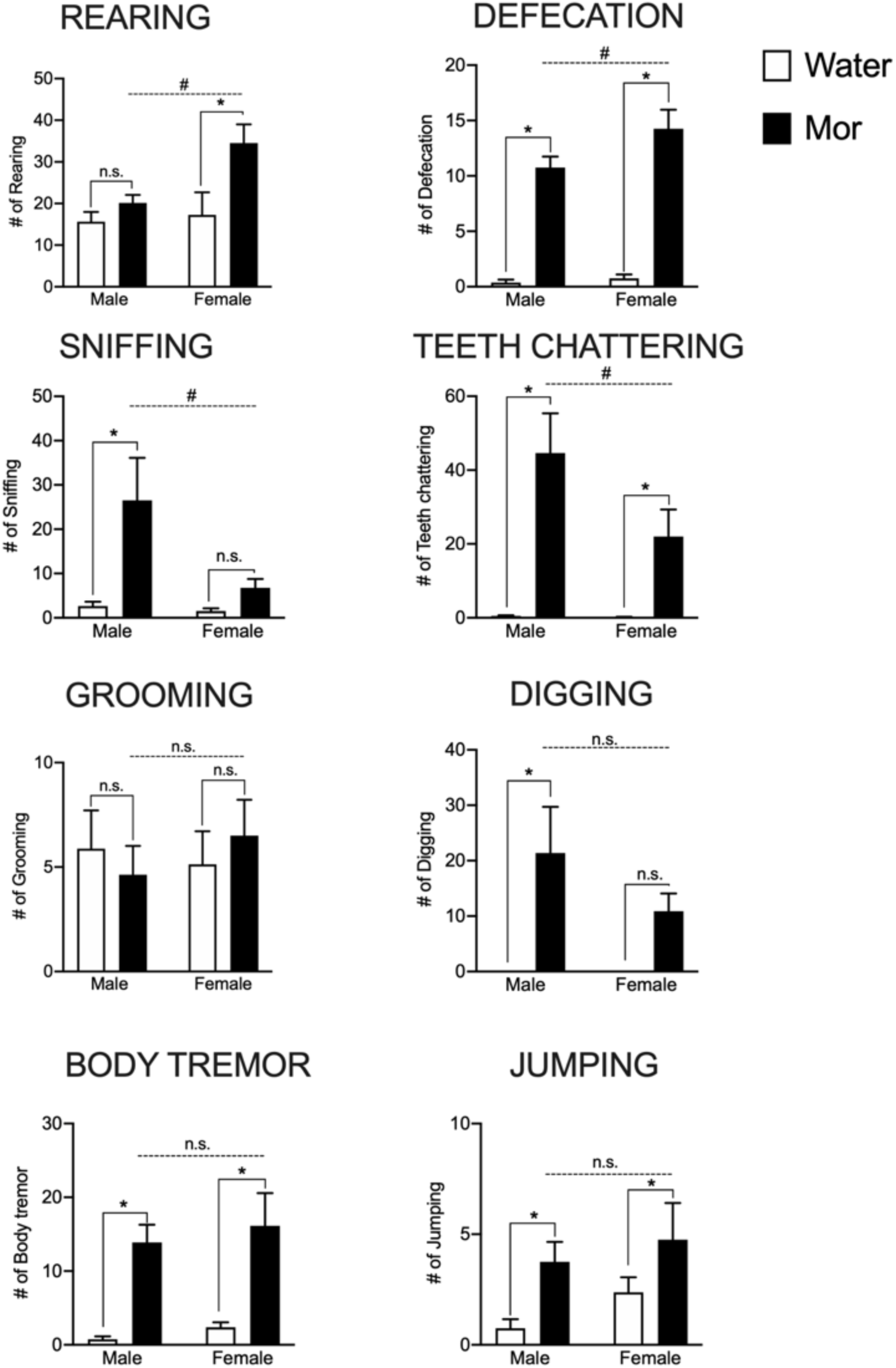
Withdrawal signs induced by naloxone injection (2 mg/kg, s.c.) in male and female morphine-dependent rats. Wistar rats treated with morphine (s.c.) for 10 days (Mor; n=8) were compared to the control group treated with distilled water (Water; n=8) in the number of withdrawal signs such as rearing, defecation, sniffing, teeth chattering, grooming, digging, body tremor and jumping. Data are expressed as mean ± SEM. * indicates p<0.05 when compared to the respective control group treated with water. # indicates p<0.05 when compared to the respective male group treated with morphine (Mor). n.s. indicates p >0.05. Two-way ANOVA followed by Bonferroni’s multiple comparison test.

**Table 1.** Values of Two-way ANOVA analysis of naloxone-precipitated withdrawal signs (treatment, sex and interaction between these factors).

| Withdrawal sign | Treatment |  | Sex |  | Interaction |  |
| --- | --- | --- | --- | --- | --- | --- |
|  | F (DFn, DFd) | p value | F (DFn, DFd) | p value | F (DFn, DFd) | p value |
| Grooming | (1, 28) = 0.001 | p = 0.9699 | (1, 28) = 0.11 | p = 0.7344 | (1, 28) = 0.63 | p = 0.4307 |
| Digging | (1, 28) = 13.04 | **p = 0.0012\*** | (1, 28) = 1.38 | p = 0.2496 | (1, 28) = 1.38 | p = 0.2496 |
| Defecation | (1, 28) = 136.2 | **p < 0.0001\*** | (1, 28) = 3.58 | p = 0.0686 | (1, 28) = 2.33 | p = 0.1378 |
| Rearing | (1, 28) = 7.98 | **p = 0.0086\*** | (1, 28) = 4.32 | **p = 0.0469\*** | (1, 28) = 2.74 | p = 0.1088 |
| Sniffing | (1, 28) = 8.65 | **p = 0.0065\*** | (1, 28) = 4.44 | **p = 0.0440\*** | (1, 28) = 3.54 | p = 0.0703 |
| Teeth Chattering | (1, 28) = 25.64 | **p < 0.0001\*** | (1, 28) = 3.11 | p = 0.0885 | (1, 28) = 2.91 | p = 0.0989 |
| Body tremor | (1, 28) = 27.61 | **p < 0.0001\*** | (1, 28) = 0.57 | p = 0.4550 | (1, 28) = 0.01 | p = 0.9036 |
| Jumping | (1, 28) = 6.84 | **p = 0.0142\*** | (1, 28) = 1.63 | p = 0.2119 | (1, 28) = 0.09 | p = 0.7632 |

Bonferroni’s multiple comparison test indicated that morphine-dependent male rats showed differences in the number of withdrawal signs of defecation (p<0.0001), sniffing (p=0.0040), teeth chattering (p<0.0001), digging (p=0.0042), body tremors (p=0.0023) and jumping (p<0.05), when compared to the water-treated male rats. No differences were detected in the withdrawal signs of rearing (p=0.83) and grooming (p>0.99) (Figure 3).

In morphine-dependent female rats, Bonferroni’s multiple comparison test revealed differences in the number of withdrawal signs of rearing (p=0.0074), defecation (p<0.0001), teeth chattering (p=0.0494), body tremors (p=0.0014) and jumping (p<0.05), when compared to the water-treated female rats. No differences were detected in the withdrawal signs of sniffing (p=0.9190), grooming (p>0.99) and digging (p=0.1922) (Figure 3).

Bonferroni’s multiple comparison test revealed sex-differences in the number of withdrawal signs of rearing (p=0.0267), defecation (p=0.0446), sniffing (p=0.0174) and teeth chattering (p=0.0411), but not grooming, digging, body tremor and jumping (p>0.05). Morphine-dependent female rats showed a higher number of defecation and rearing, and a lower number of sniffing and teeth chattering, than morphine-dependent male rats (Figure 3).

Table 2 shows the estimate percentages for each withdrawal sign calculated from the total number of withdrawal signs elicited during the 30-minutes period of the test in morphine dependent male and female rats. The sign of teeth chattering was the most preeminent manifestation of opioid-withdrawal for male morphine-dependent rats (29%), whereas the sign of rearing was the most preeminent for female rats (31%). Furthermore, student’s T test indicated that female morphine-dependent rats showed a higher percentage of rearing [t=4.105; df=14; p=0.0011] and defecation signs [t=2.248; df=14; p=0.0412] than male morphine-dependent rats. No differences were found between male and female dependent rats in the estimate mean percentages of grooming [t=1.131; df=14; p=0.2771], digging [t=0.9333; df=14; p=0.3670], sniffing [t=1.754; df=14; p=0.1013], teeth chattering [t=1.250; df=14; p=0.2316], body tremor [t=0.7062; df=14; p=0.4917] and jumping [t=0.8120; df=14; p=0.4304].

**Table 2.** Estimate mean percentage of each withdrawal sign from the total number of withdrawal signs during the test period (morphine-dependent group only).

| Withdrawal sign | Male | Female | p value |
| --- | --- | --- | --- |
| Grooming | 3.68 ± 1.16 | 5.67 ± 1.31 | 0.2771 |
| Rearing | 14.68 ± 1.72 | 30.53 ± 3.46 | **0.0011\*** |
| Defecation | 8.18 ± 1.18 | 12.60 ± 1.57 | **0.0412\*** |
| Digging | 14.15 ± 4.29 | 9.38 ± 2.92 | 0.3670 |
| Sniffing | 17.20 ± 6.12 | 5.95 ± 1.91 | 0.1013 |
| Teeth chattering | 29.04 ± 4.84 | 19.15 ± 6.25 | 0.2316 |
| Body tremor | 10.24 ± 1.76 | 12.52 ± 2.70 | 0.4917 |
| Jumping | 2.83 ± 0.77 | 4.28 ± 1.60 | 0.4304 |

On the next set of experiments, both male and female rats were treated again with different doses of CBD (3, 10 and 30 mg/kg, i.p.), 10 minutes before the injection of morphine in the morning of the 11^th^ day during the dependence protocol. As shown in Figure 4, One-way ANOVA indicated significant effect of treatment with CBD (day 11) in the total number of the withdrawal signs of sniffing [F(3, 35) = 4.785; p= 0.0067] and teeth chattering [F(3, 35) = 4.702; p= 0.0073] precipitated by naloxone injection in male morphine-dependent rats. Bonferroni’s multiple comparison test revealed that treatment with CBD with the doses of 3, 10 and 30 mg/kg (CBD 3 – Mor, CBD 10 – Mor and CBD 30 – Mor), reduced the total number of sniffing and teeth chattering Although One-way ANOVA did not indicate effect of treatment in the total number of body tremors in male rats, Bonferroni’s multiple comparison test revealed significant effect of treatment with 10 mg/kg of CBD (CBD 10 – Mor; p= 0.0454) (Figure 4).

**Figure 4.**
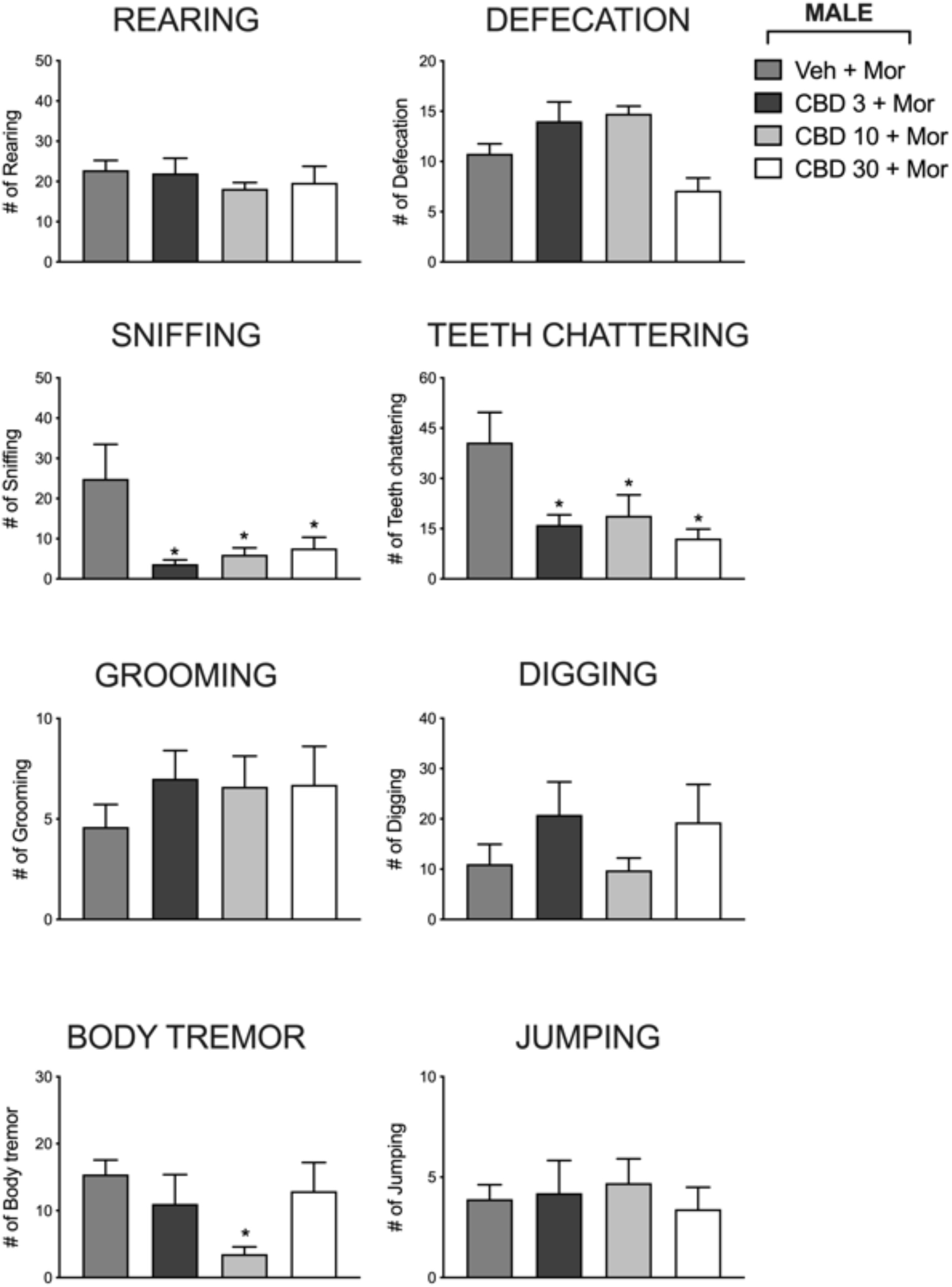
Effect of CBD treatment in the withdrawal signs rearing, defecation, sniffing, teeth chattering, grooming, digging, body tremor and jumping in male morphine-dependent rats. Groups of male Wistar rats treated with CBD (3, 10 or 30 mg/kg; i.p.) before every injection of morphine (s.c.) for 10 days were compared to the control group treated with vehicle (Veh-Mor). Data are expressed as mean ± SEM, n=8-10 per group. * indicates p<0.05. One-way ANOVA followed by Bonferroni’s multiple comparison test.

Furthermore, one-way ANOVA indicated significant effect of treatment with CBD in the withdrawal of defecation [F(3, 32) = 7.607; p= 0.0454]. However, Bonferroni’s multiple comparison test did not evidence significant effect of CBD in the total number of defecations in any of the doses tested (p>0.05). No significant differences were observed in the total number of the withdrawal signs of rearing [F(3, 36) = 0.4518; p= 0.7176], grooming [F(3, 36) = 0.5210; p= 0.6706], digging [F(3, 34) = 0.9698; p= 0.4183] and jumping [F(3, 36) = 0.2027; p= 0.8939], in the groups treated with CBD (CBD 3 – Mor, CBD 10 – Mor and CBD 30 – Mor), when compared to the control group (Veh-Mor) in male rats (Figure 4).

As shown in Figure 5, CBD treatment had no effect in the withdrawal signs of female rats, in any of the doses tested (CBD 3 – Mor, CBD 10 – Mor and CBD 30 – Mor). One-way ANOVA indicated no significant effect of treatment in the total number of withdrawal signs of rearing [F(3, 36) = 0.5418; p= 0.6568], defecation [F(3, 36) = 1.616; p= 0.2026], sniffing [F(3, 36) = 0.4368; p= 0.7280], teeth chattering [F(3, 36) = 1.019; p= 0.3955], grooming [F(3, 36) = 0.1789; p= 0.9100], digging [F(3, 36) = 1.965; p= 0.1366], body tremor [F(3, 36) = 0.1517; p= 0.9240] and jumping [F(3, 36) = 1.744; p= 0.1755] in female morphine-dependent Wistar rats, when compared to the control group (Veh-Mor).

**Figure 5.**
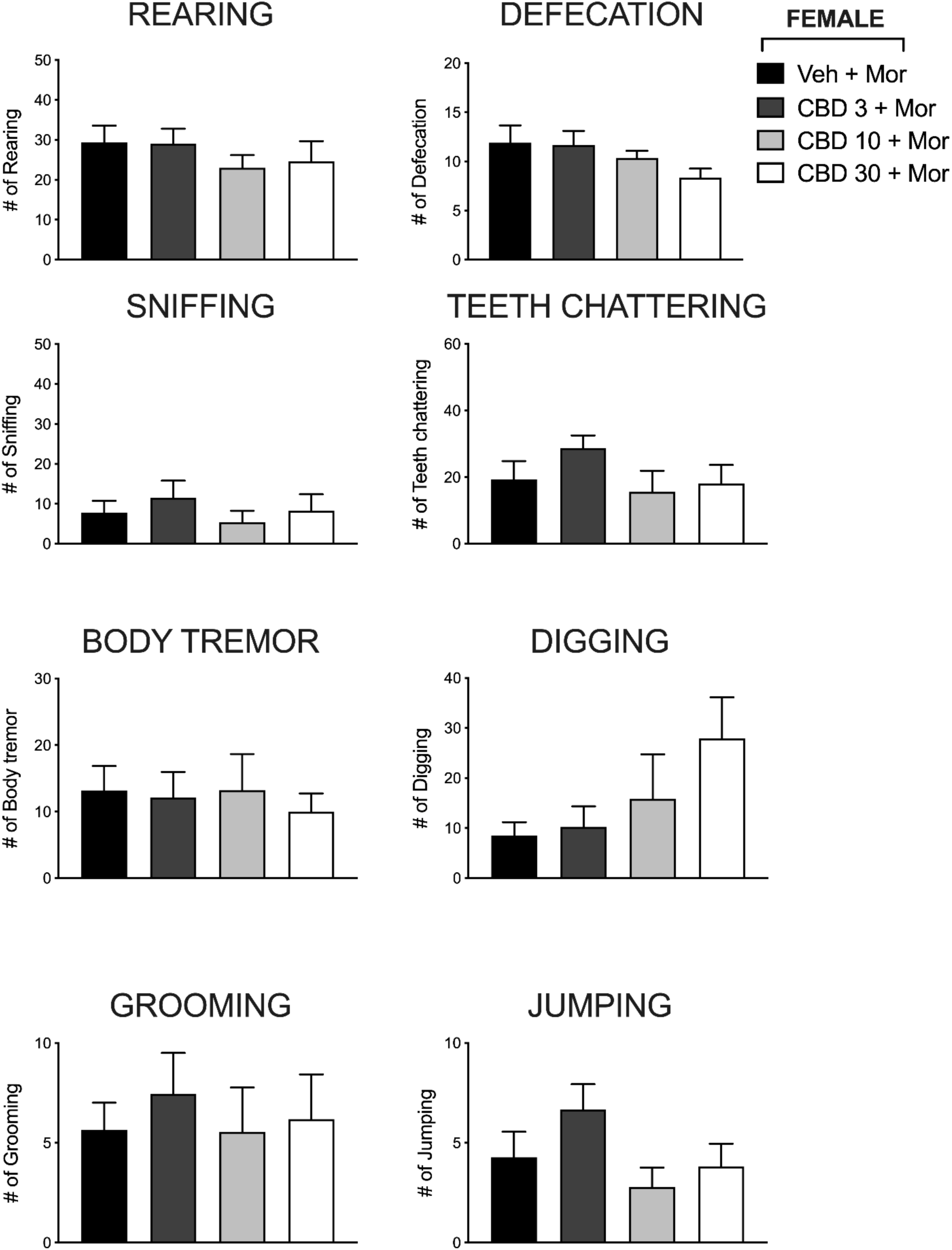
Effect of CBD treatment in the withdrawal signs of rearing, defecation, sniffing, teeth chattering, grooming, digging, body tremor and jumping in female morphine-dependent rats. Groups of female Wistar rats treated with CBD (3, 10 or 30 mg/kg; i.p.) before every injection of morphine (s.c.) for 10 days were compared to the control group treated with vehicle (Veh-Mor). Data are expressed as mean ± SEM, n=9-11 per group. * indicates p<0.05. One-way ANOVA followed by Bonferroni’s multiple comparison test.

## 4. Discussion

The primary finding of our study is that morphine sulfate solution, prepared through filtration of tablets and given subcutaneously twice daily for 10 days, induced thermal hyperalgesia and physical dependence in both male and female rats, as shown by reduced latency in the hot plate test and the different withdrawal signs precipitated by naloxone injection, respectively. In addition, our study shows evidence of sex-differences in the expression of withdrawal signs in Wistar rats. Lastly, the present study demonstrated that CBD reduces morphine induced hyperalgesia in both sexes and attenuates withdrawal signs, such sniffing, teeth chattering and body tremor, in morphine-dependent male rats, but not female rats.

Morphine-induced physical dependence and hyperalgesia in rodents can be induced by different methods such as implantation of subcutaneous pellets [44,45], oral ingestion of morphine [46] and subcutaneous daily injections [39], as well as different dose ranges [39,47]. The solution made by a filtration of crushed tablets may not represent the first choice of solution for the investigation of the addictive effects of morphine in a non-clinical study, due to the potential presence of excipients. In addition, further studies investigating potential effect differences between filtered solution and pure morphine solution are necessary, and for that reason might indicate a limiting factor in our study. Nonetheless, solutions made of commercial tablets is one of the alternatives applied by users as a source of drug, and it represents a feature encountered in the clinical setting[36,37].

In our study male and female rats treated for 10 days with morphine solution prepared by filtration of commercial tablets developed thermal hyperalgesia starting 7 days after the beginning of treatments. This result is in accordance with previous studies showing that chronic exposure to morphine induces thermal hyperalgesia in non-clinical models [40,48]. There is a volume of conflicting data indicating that the hyperalgesic effect of morphine might be different between sexes. For instance, Holtman and Wala [49] showed that female rats had a markedly greater response of thermal hyperalgesia to a low dose of morphine, compared to male rats. o et al. [50] demonstrated that low dose infusion of morphine (1.6 mg/kg/day) caused hyperalgesia that was evident earlier in female mice, when compared to male mice, and this manifestation dissipated earlier in males than in females. However, a larger dose of morphine (40 mg/kg/day) induced hyperalgesia that was identical in onset, magnitude, and duration between sexes. The impact of the estrous cycle on morphine-induced hyperalgesia is not well defined in rodents [51], but it has been reported that ovariectomy causes hyperalgesia, induced by low doses of morphine in female mice, to dissipate in a manner similar to males, an effect blocked by estrogen treatment [50], which indicates that ovarian steroids might divert the hyperalgesic mechanisms of morphine. Thus, further studies are required for the evaluation of potential fluctuations in the estrous cycle of female rats and their impact on the morphine-induced hyperalgesia protocol used in the current study.

Morphine-induced hyperalgesia involves multiple pathways, but studies have reported an important role of TRPV1 receptors in the development of this manifestation. For instance, Vardanyan and colleagues [45] reported that mice with morphine pellets implanted subcutaneously developed thermal hyperalgesia 7 days after pellet implantation. Furthermore, this same study has reported that thermal hyperalgesia was mediated by an increase in TRPV1 receptor function. In our study, repeated treatment with CBD (30 mg/kg) reduced thermal hyperalgesia in morphine-dependent male and female rats. Accordingly, CBD has shown potential to treat different experimental conditions of pain [43,52]. In addition, studies have indicated that co-treatment with TRPV1 receptor antagonist was able to block the anti-hyperalgesic effect of CBD, suggesting that CBD may suppress pain through sensitization of TRPV1 receptors [53,54]. Altogether, these results indicate that CBD reduces morphine-induced hyperalgesia, and this effect might be potentially mediated by TRPV1 receptor activation. Further studies are necessary to evaluate the role of TRPV1 receptors in the effect of CBD on OIH in male and female rodents.

Other than hyperalgesia, withdrawal syndrome is also a complication of opioid use and may involve unpleasant and aversive symptoms (i.e., tachycardia, agitation, anxiety) [55,56], these symptoms might lead to opioid craving and relapse in patients [57]. In the present study, both morphine-dependent male and female rats demonstrated significant withdrawal signs (rearing, grooming, digging, defecation, teeth chattering, sniffing, body tremor and jumping) after naloxone injection, which is accordance to other non-clinical reports [39,47]. Precipitated withdrawal behavior can be expressed in humans and laboratory animals as affective and emotional symptoms, as well as somatic signs. A spectrum of behavior occurs in both rats and mice in a similar manner, as shown by only few studies that investigated potential differences between species[58]. Studies have also reported that withdrawal behavior can vary from protocols investigating precipitated or spontaneous withdrawal[59]. Evaluating animals in observation chambers, for instance, is one factor that can play a role in the differences in withdrawal behavior, as space constraint might limit the number of wet dog shakes, abdominal constrictions and writhing behavior observed during a determined time of testing [58]. In addition, jumping behavior might be impacted by the height of observation chambers, as rats in withdrawal have shown to jump less as the height of the chamber increases [60]. Due to these conditions, behaviors like teeth chattering, salivation, sniffing, hyperirritability, and others, found in the current study, might be observed more often.

Our data indicated that female and male rats showed discrepancies in the total number of withdrawal signs manifested during the test. Morphine-dependent female rats preeminently showed the withdrawal signs of rearing and defecation, whereas male rats showed preeminently the signs of sniffing and teeth chattering. Multiple studies indicate sex-differences in the behavioral strategies of rodent’s manifestation of behavior [61]. For instance, in fear conditioning behavior, female rodents are more likely to exhibit alternate responses to fear, such as escape-like darting response, rather than freezing, which is expressed in male rodents [62]. Regarding addiction, during precipitated nicotine withdrawal, more somatic withdrawal signs (body shakes, head shakes, ptosis, and others) were found in male rats, in comparison to female rats [63]. However, studies exploiting sex differences in opioid withdrawal are lacking, and the few available show inconsistent results that report greater sensitivity of male rodents or female rodents in comparison to each other, or even no sex differences [64].

There is increasing evidence of interaction between the opioidergic, and endocannabinoid systems and modulation of cannabinoid receptors has shown to exert effects on the rewarding effects of opioids [65]. For instance, the administration of a cannabinoid antagonist is reported to attenuate morphine-induced CPP, whereas administration of opioid antagonist blocks cannabinoid-induced CPP [19,20]. These mechanisms are not well explored in humans, but a study has shown that individuals who use opioids have increased cannabinoid type 1 (CB_1_) receptor expression in opioid rewarding pathways [66]. Regarding opioid withdrawal, exogenous endocannabinoids (2-arachidonoylglycerol and tetrahydrocannabinol) seem to exert relief of withdrawal symptoms in rodents (paw tremors, diarrhea, jumps and others) [67,68].

In the last decade, CBD, a non-psychotomimetic constituent in the *Cannabis* plant, has gained popularity in the medical community. CBD is the second most common phytocannabinoid, does not directly activate CB_1_ or CB_2_ receptors [69] and it lacks the rewarding properties which is inherent in another phytocannabinoid, Delta 9-Tetrahydrocannabinol (THC) [31,70]. Further experiments are necessary to investigate if CBD treatment alone, given in the schedule proposed in our study, could potentially induce withdrawal signs. Nonetheless, studies have indicated that CBD treatment does not induce hedonic effects [70] or conditioned place preference by itself, and does not produce withdrawal behavior, such as grooming, rubbing and rearing, in mice [71]. A randomized clinical trial has also observed that abrupt interruption of a short-term treatment with CBD did not induce withdrawal syndrome [72], which is in accordance with animal studies showing that CBD displays in rodents a comparable motivation to the consumption of water, for instance [71]. In addition, CBD displays a wide range of therapeutic effects in different medical and psychological conditions, such as substance use disorder [73].

Recently, a double-blind randomized placebo-controlled trial by Hurd and coworkers [32] demonstrated a promising and safe effect of CBD on drug-cue-induced craving in abstinent individuals with heroin use disorder. In comparison to placebo, CBD treatment reduced cue-induced craving, heart rate and salivary cortisol levels. In addition, no serious adverse effects were reported in the trial. In non-clinical models of opioid addiction, CBD has shown efficacy in reducing cue-induced heroin seeking behavior in rats, in both short (24 hours) and long (2 weeks) periods after administration [35]. In the conditioned place preference (CPP) paradigm, co-treatment with CBD reduced morphine induced CPP in mice [34] and rats [74]. Moreover, CBD treatment blocked conditioned place aversion induced by naloxone injection in rats [74]. In our study, an acute treatment with CBD (doses of 3, 10 and 30 mg/kg) 10 minutes before the morphine injection on day 11 of the protocol, attenuated the significant frequent withdrawal signs of sniffing and teeth chattering, and the withdrawal sign of body tremors (dose of CBD 10 mg/kg) in morphine-dependent male rats that received an administration of naloxone. Not surprisingly, these results corroborate with other studies showing that CBD may produce U-shaped dose response curves of effect in behavior tests with rodents [75,76] and in clinical trials evaluating its anxiolytic properties [77]. In oxycodone-precipitated and spontaneous withdrawal, CBD induced significant reduction of gastrointestinal symptoms (fecal boli count), in both male and female mice[59]. Lastly, in accordance with our results, CBD has shown potential to reduce somatic withdrawal signs and mechanical hyperalgesia in rats during acute and protracted abstinence of another addictive drug such as nicotine [78].

CBD is well known for its wide range of cellular mechanisms [79]. However, studies have tried to narrow possible mechanisms for CBD’s action on opioid-induced addictive behaviors. For instance, treatment with CBD has shown to provide normalization of AMPA (α-amino-3-hydroxy-5-methyl-4-isoxazolepropionic acid) GluR1 and cannabinoid type-1 receptor (CB1) expression in nucleus accumbens, that was once increased by cue-induced heroin-seeking in rats [35]. Furthermore, CBD treatment inhibited the reward-facilitating effect of morphine evaluated in the intracranial self-stimulation paradigm. This effect was blocked by a selective 5-HT1A antagonist injection in the dorsal raphe nucleus, suggesting a serotonergic mechanism involved in the effect of CBD on the rewarding effects of opioids[31]. Altogether these results reveal the ability of CBD to interfere in the rewarding effects of opioids and in the severity of withdrawal signs found in the morphine-dependent male rats.

In our study, CBD significantly reduced withdrawal signs in male rats, but did not produce attenuation of withdrawal signs in morphine-dependent female rats. As stated previously, non-clinical studies have demonstrated the higher rewarding effects of opioid exposure in female rats. Regarding the rewarding effects of morphine itself, Cicero *et al*. [80] has demonstrated that both male and female rats show the rewarding effects of morphine through increased conditioned place preference, but only females continue to show this effect in doses up to 30 mg/kg. Moreover, the oral consumption of water containing morphine was higher in female rats, in comparison to male rats [46]. Intravenous self-administration of opioids, such as morphine and heroin, was also reported to be greater in female rats than in males in an operant conditioning paradigm [81]. Although the impact of gonadal hormones was not approached in our protocols, studies have reported that the presence of estradiol is involved in the development and in the augmentation of addictive phenotypes in female rats treated with cocaine or morphine [82–84], which suggests that intact females are reliable to reproduce addictive behaviors after long-term exposure to opioids.

Studies have revealed contrasting results of sex differences in the effects of cannabinoids. For instance, cannabinoids have shown to produce greater antinociception, catalepsy, sexual behavior, and anxiety in female, when compared to male rats [85], but locomotor and thermoregulatory responses remain similar between both sexes [86]. Furthermore, cannabinoid self-administration has been reported significant higher in intact female rats, than in male rats, whereas ovariectomized females were less sensitive to the reinforcing effects of cannabinoids [87]. Overall, there is limited literature on the sex differences in the effect of CBD on addictive behavior, and further studies are required to evaluate the mechanisms involved in the sex dependent effect of CBD in the withdrawal signs of morphine-dependent rats.

## 5. Conclusions

Taken together, our findings point to the potential effect of CBD as a treatment for OIH and withdrawal syndrome in experimental morphine dependence. Our results have revealed the efficacy of CBD in the treatment of OIH in both male and female rats, as well as differences in the withdrawal signs precipitated by naloxone injection in male and female morphine-dependent rats. Importantly, morphine-dependent male and female rats respond different to a treatment with CBD, which suggests a sex-difference in the pharmacological effect of this phytocannabinoid in addictive behaviors. Further studies are necessary to clarify the mechanisms involved in the sex differences observed in the response to CBD treatment in male and female rodents expressing opioid withdrawal behavior. Nonetheless, CBD might represent a potential therapeutic target to treat complications of OUD.

## Author Contribution

**C.H.A.J, J.V.**: Conceptualization, Formal Analysis, Investigation, Methodology, Project Administration, Supervision, Visualization, Data Curation, Writing – original draft, Writing – review and editing. **B.B.S., M.A.R.B., M.V.F.**: Conceptualization, Formal Analysis, Investigation, Methodology and Data Curation. **F.F.:** Investigation, Methodology and Data Curation. **K.G., A.S.C., D.E.P.S.:** Funding acquisition, Visualization, Writing – review and editing. **J.M.C.:** Funding acquisition, Visualization, Project Administration, Supervision, Writing – review and editing.

## Aknowledgements

CHAJ, JV, MVF, BBS and MARB are recipients of Coordenação de Aperfeiçoamento de Pessoal de Nível Superior (CAPES) fellowships (Finance Code 001). We thank BSPG-Pharm (Sandwich, UK) who kindly donated CBD. The funders had no role in the design and conduct of the study; collection, management, analysis, and interpretation of the data; preparation, review, or approval of the manuscript; and decision to submit the manuscript for publication. All authors read and approved the final manuscript.

## List of Abbreviations

5HT1A: serotonin 1A receptor
AMPA: α-amino-3-hydroxy-5-methyl-4-isoxazolepropionic acid
ANOVA: analysis of variance
CBD: cannabidiol
CB_1_: cannabinoid receptor type 1
CPP: conditioned place preference
MOR: morphine
OUD: opioid use disorder
OIH: opioid-induced hyperalgesia
TRPV1: transient receptor potential channel subfamily V member 1
Veh: vehicle;

## References

1. Dydyk AM, Jain NK, Gupta M. Opioid Use Disorder. StatPearls. Published online July 12, 2021.

2. Buresh M, Stern R, Rastegar D. Treatment of opioid use disorder in primary care. BMJ. 2021;373. doi:10.1136/BMJ.N784

3. Olfson M, Rossen LM, Wall MM, Houry D, Blanco C. Trends in Intentional and Unintentional Opioid Overdose Deaths in the United States, 2000-2017. JAMA. 2019;322(23):2340–2342. doi:10.1001/JAMA.2019.16566

4. WHO. Opioid overdose. Opioid overdose, World Health Organization. Published 2021. Accessed December 15, 2021. https://www.who.int/news-room/fact-sheets/detail/opioid-overdose

5. Roth ME, Cosgrove KP, Carroll ME. Sex differences in the vulnerability to drug abuse: a review of preclinical studies. Neurosci Biobehav Rev. 2004;28(6):533–546. doi:10.1016/J.NEUBIOREV.2004.08.001

6. Becker JB, McClellan ML, Reed BG. Sex differences, gender and addiction. J Neurosci Res. 2017;95(1-2):136. doi:10.1002/JNR.23963

7. Robinson HL, Banks ML. Adding dopamine to the complexity of sex differences in opioid reinforcement. Neuropsychopharmacology 2021 46:*10*. 2021;46(10):1705-1706. doi:10.1038/s41386-021-01060-z

8. Huhn AS, Tompkins DA, Campbell CM, Dunn KE. Individuals with Chronic Pain Who Misuse Prescription Opioids Report Sex-Based Differences in Pain and Opioid Withdrawal. Pain Med. 2019;20(10):1942–1947. doi:10.1093/PM/PNY295

9. Bobzean SAM, Kokane SS, Butler BD, Perrotti LI. Sex differences in the expression of morphine withdrawal symptoms and associated activity in the tail of the ventral tegmental area. Neurosci Lett. 2019;705:124. doi:10.1016/J.NEULET.2019.04.057

10. Huhn AS, Berry MS, Dunn KE. Review: Sex-based Differences in Treatment Outcomes for Persons with Opioid Use Disorder. Am J Addict. 2019;28(4):246. doi:10.1111/AJAD.12921

11. Enman NM, Reyes BAS, Shi Y, Valentino RJ, Van Bockstaele EJ. Sex differences in morphine-induced trafficking of mu-opioid and corticotropin-releasing factor receptors in locus coeruleus neurons. Brain Res. 2019;1706:75–85. doi:10.1016/J.BRAINRES.2018.11.001

12. Zhou J, Ma R, Jin Y, et al. Molecular mechanisms of opioid tolerance: From opioid receptors to inflammatory mediators (Review). Exp Ther Med. 2021;22(3). doi:10.3892/ETM.2021.10437

13. Lowl Y, Clarke2 CF, Huhl BK. Opioid-induced hyperalgesia: a review of epidemiology, mechanisms and management. IReview Article Singapore Med J. 2012;53(5):357–360.

14. Wilson SH, Hellman KM, James D, Adler AC, Chandrakantan A. Mechanisms, diagnosis, prevention and management of perioperative opioid-induced hyperalgesia. Pain Manag. 2021;11(4):405–417. doi:10.2217/PMT-2020-0105

15. Doverty M, White JM, Somogyi AA, Bochner F, Ali R, Ling W. Hyperalgesic responses in methadone maintenance patients. Pain. 2001;90(1-2):91–96. doi:10.1016/S0304-3959(00)00391-2

16. Comer S, Cunningham C, Fishman MJ, et al. National Practice Guideline for the Use of Medications in the Treatment of Addiction Involving Opioid Use ASAM.; 2015.

17. Srivastava AB, Mariani JJ, Levin FR. New directions in the treatment of opioid withdrawal. The Lancet. 2020;395(10241):1938–1948. doi:10.1016/S0140-6736(20)30852-7

18. Graczyk M, Łukowicz M, Dzierzanowski T. Prospects for the Use of Cannabinoids in Psychiatric Disorders. Front Psychiatry. 2021;12:276. doi:10.3389/FPSYT.2021.620073/BIBTEX

19. Singh ME, Verty ANA, McGregor IS, Mallet PE. A cannabinoid receptor antagonist attenuates conditioned place preference but not behavioural sensitization to morphine. Brain Res. 2004;1026(2):244–253. doi:10.1016/J.BRAINRES.2004.08.027

20. Braida D, Iosuè S, Pegorini S, Sala M. Δ9-Tetrahydrocannabinol-induced conditioned place preference and intracerebroventricular self-administration in rats. Eur J Pharmacol. 2004;506(1):63–69. doi:10.1016/J.EJPHAR.2004.10.043

21. Gossop M, Battersby M, Strang J. Self-detoxification by opiate addicts. A preliminary investigation. Br J Psychiatry. 1991;159(AUG.):208–212. doi:10.1192/BJP.159.2.208

22. Epstein DH, Preston KL. No Evidence for Reduction of Opioid-Withdrawal Symptoms by Cannabis Smoking During a Methadone Dose Taper. Am J Addict. 2015;24(4):323. doi:10.1111/AJAD.12183

23. Fattore L, Deiana S, Spano SM, et al. Endocannabinoid system and opioid addiction: Behavioural aspects. Pharmacol Biochem Behav. 2005;81(2):343–359. doi:10.1016/J.PBB.2005.01.031

24. Atalay S, Jarocka-Karpowicz I, Skrzydlewska E. Antioxidative and Anti-Inflammatory Properties of Cannabidiol. Antioxidants. 2020;9(1). doi:10.3390/ANTIOX9010021

25. Lim K, See YM, Lee J. A Systematic Review of the Effectiveness of Medical Cannabis for Psychiatric, Movement and Neurodegenerative Disorders. Clinical Psychopharmacology and Neuroscience. 2017;15(4):301. doi:10.9758/CPN.2017.15.4.301

26. Chaves YC, Genaro K, Crippa JA, da Cunha JM, Zanoveli JM. Cannabidiol induces antidepressant and anxiolytic-like effects in experimental type-1 diabetic animals by multiple sites of action. Metab Brain Dis. 2021;36(4):639–652. doi:10.1007/S11011-020-00667-3/TABLES/2

27. Viudez-Martínez A, García-Gutiérrez MS, Navarrón CM, et al. Cannabidiol reduces ethanol consumption, motivation and relapse in mice. Addiction biology. 2018;23(1):154–164. doi:10.1111/ADB.12495

28. Castillo-Arellano J, Canseco-Alba A, Cutler SJ, León F. The Polypharmacological Effects of Cannabidiol. Molecules. 2023;28(7). doi:10.3390/MOLECULES28073271

29. Galaj E, Bi GH, Yang HJ, Xi ZX. Cannabidiol attenuates the rewarding effects of cocaine in rats by CB2, 5-HT1A and TRPV1 receptor mechanisms. Neuropharmacology. 2020;167. doi:10.1016/J.NEUROPHARM.2019.107740

30. Luján MÁ, Castro-Zavala A, Alegre-Zurano L, Valverde O. Repeated Cannabidiol treatment reduces cocaine intake and modulates neural proliferation and CB1R expression in the mouse hippocampus. Neuropharmacology. 2018;143:163–175. doi:10.1016/J.NEUROPHARM.2018.09.043

31. Katsidoni V, Anagnostou I, Panagis G. Cannabidiol inhibits the reward-facilitating effect of morphine: involvement of 5-HT 1A receptors in the dorsal raphe nucleus. Addiction Biology. 2012;18(2):286–296. doi:10.1111/j.1369-1600.2012.00483.x

32. Hurd YL, Spriggs S, Alishayev J, et al. Cannabidiol for the reduction of cue-induced craving and anxiety in drug-abstinent individuals with heroin use disorder: A double-blind randomized placebo-controlled trial. American Journal of Psychiatry. 2019;176(11):911–922. doi:10.1176/appi.ajp.2019.18101191

33. Bhargava HN. Effect of some cannabinoids on naloxone-precipitated abstinence in morphine-dependent mice. Psychopharmacology 1976 49:*3*. 1976;49(3):267-270. doi:10.1007/BF00426828

34. Markos JR, Harris HM, Gul W, Elsohly MA, Sufka KJ. Effects of Cannabidiol on Morphine Conditioned Place Preference in Mice. Planta Med. 2018;84(4):221–224. doi:10.1055/s-0043-117838

35. Ren Y, Whittard J, Higuera-Matas A, Morris C V., Hurd YL. Cannabidiol, a Nonpsychotropic Component of Cannabis, Inhibits Cue-Induced Heroin Seeking and Normalizes Discrete Mesolimbic Neuronal Disturbances. The Journal of Neuroscience. 2009;29(47):14764. doi:10.1523/JNEUROSCI.4291-09.2009

36. McLean S, Bruno R, Brandon S, de Graaff B. Effect of filtration on morphine and particle content of injections prepared from slow-release oral morphine tablets. Harm Reduct J. 2009;6(1):1–13. doi:10.1186/1477-7517-6-37/FIGURES/7

37. Keijzer L. Reducing harm through the development of good preparation practices for the injection of slow release morphine sulphate capsules. Harm Reduct J. 2020;17(1):1–9. doi:10.1186/S12954-020-00389-W/TABLES/1

38. National Center for Biotechnology Information. PubChem Compound Summary for CID 5288826, Morphine. Published 2021. https://pubchem.ncbi.nlm.nih.gov/compound/Morphine.

39. Zamani N, Hassanian-Moghaddam H, Bayat A, et al. Reversal of opioid overdose syndrome in morphine-dependent rats using buprenorphine. Toxicol Lett. 2015;232(3):590–594. doi:10.1016/j.toxlet.2014.12.007

40. Datta U, Kelley LK, Middleton JW, Gilpin NW. Positive allosteric modulation of the cannabinoid type-1 receptor (CB1R) in periaqueductal gray (PAG) antagonizes anti-nociceptive and cellular effects of a mu-opioid receptor agonist in morphine-withdrawn rats. Psychopharmacology (Berl). 2020;237(12):3729–3739. doi:10.1007/S00213-020-05650-5

41. Pacheco SDG, Gasparin AT, Jesus CHA, et al. Antinociceptive and Anti-Inflammatory Effects of Bixin, a Carotenoid Extracted from the Seeds of Bixa orellana. Planta Med. 2019;85(16):1216–1224.

42. Genaro K, Fabris D, Arantes ALF, Zuardi AW, Crippa JAS, Prado WA. Cannabidiol Is a Potential Therapeutic for the Affective-Motivational Dimension of Incision Pain in Rats. Front Pharmacol. 2017;8:391. doi:10.3389/fphar.2017.00391

43. Jesus CHA, Redivo DDB, Gasparin AT, et al. Cannabidiol attenuates mechanical allodynia in streptozotocin-induced diabetic rats via serotonergic system activation through 5-HT1A receptors. Brain Res. 2019;1715:156–164. doi:10.1016/J.BRAINRES.2019.03.014

44. Wang F, Meng J, Zhang L, Johnson T, Chen C, Roy S. Morphine induces changes in the gut microbiome and metabolome in a morphine dependence model. Scientific Reports 2018 8:*1*. 2018;8(1):1-15. doi:10.1038/s41598-018-21915-8

45. Vardanyan A, Wang R, Vanderah TW, et al. TRPV1 receptor in expression of opioid-induced hyperalgesia. J Pain. 2009;10(3):243–252. doi:10.1016/J.JPAIN.2008.07.004

46. Alexander BK, Coambs RB, Hadaway PF. The effect of housing and gender on morphine self-administration in rats. Psychopharmacology 1978 58:*2*. 1978;58(2):175-179. doi:10.1007/BF00426903

47. Rezaei Z, Kourosh-Arami M, Azizi H, Semnanian S. Orexin type-1 receptor inhibition in the rat lateral paragigantocellularis nucleus attenuates development of morphine dependence. Neurosci Lett. 2020;724. doi:10.1016/j.neulet.2020.134875

48. Ferrini F, Lorenzo LE, Godin AG, Quang M Le, De Koninck Y. Enhancing KCC2 function counteracts morphine-induced hyperalgesia. Scientific Reports 2017 7:*1*. 2017;7(1):1-8. doi:10.1038/s41598-017-04209-3

49. Holtman JR, Wala EP. Characterization of morphine-induced hyperalgesia in male and female rats. Pain. 2005;114(1-2):62–70. doi:10.1016/J.PAIN.2004.11.014

50. Juni A, Klein G, Kowalczyk B, Ragnauth A, Kest B. Sex differences in hyperalgesia during morphine infusion: effect of gonadectomy and estrogen treatment. Neuropharmacology. 2008;54(8):1264–1270. doi:10.1016/J.NEUROPHARM.2008.04.004

51. Bodnar RJ, Kest B. Sex differences in opioid analgesia, hyperalgesia, tolerance and withdrawal: Central mechanisms of action and roles of gonadal hormones. Horm Behav. 2010;58(1):72–81. doi:10.1016/J.YHBEH.2009.09.012

52. Ward SJ, McAllister SD, Kawamura R, Murase R, Neelakantan H, Walker EA. Cannabidiol inhibits paclitaxel-induced neuropathic pain through 5-HT 1A receptors without diminishing nervous system function or chemotherapy efficacy. Br J Pharmacol. 2014;171(3):636–645. doi:10.1111/bph.12439

53. Costa B, Giagnoni G, Franke C, Trovato AE, Colleoni M. Vanilloid TRPV1 receptor mediates the antihyperalgesic effect of the nonpsychoactive cannabinoid, cannabidiol, in a rat model of acute inflammation. Br J Pharmacol. 2004;143(2):247–250. doi:10.1038/sj.bjp.0705920

54. Costa B, Trovato AE, Comelli F, Giagnoni G, Colleoni M. The non-psychoactive cannabis constituent cannabidiol is an orally effective therapeutic agent in rat chronic inflammatory and neuropathic pain. Eur J Pharmacol. 2007;556(1-3):75–83. doi:10.1016/j.ejphar.2006.11.006

55. Shah M, Huecker MR. Opioid Withdrawal. *Challenging Cases and Complication Management in Pain Medicine*. Published online October 11, 2021:15–20.

56. American Addiction Centers. Opioid Withdrawal: Signs, Symptoms & Addiction Treatment. Published online 2021.

57. Bruneau A, Frimerman L, Verner M, et al. Day-to-day opioid withdrawal symptoms, psychological distress, and opioid craving in patients with chronic pain prescribed opioid therapy. Drug Alcohol Depend. 2021;225:108787. doi:10.1016/J.DRUGALCDEP.2021.108787

58. Uddin O, Jenne C, Fox ME, Arakawa K, Keller A, Cramer N. Divergent profiles of fentanyl withdrawal and associated pain in mice and rats. bioRxiv. Published online November 16, 2020:2020.11.16.384818. doi:10.1101/2020.11.16.384818

59. Scicluna RL, Wilson BB, Thelaus SH, Arnold JC, McGregor IS, Bowen MT. Cannabidiol Reduced the Severity of Gastrointestinal Symptoms of Opioid Withdrawal in Male and Female Mice. https://home.liebertpub.com/can. Published online December 27, 2022. doi:10.1089/CAN.2022.0036

60. Azizi H, Ranjbar-Slamloo Y, Semnanian S. Height-dependent difference in the expression of naloxone-induced withdrawal jumping behavior in morphine dependent rats. Neurosci Lett. 2012;515(2):174–176. doi:10.1016/J.NEULET.2012.03.047

61. Shansky RM, Murphy AZ. Considering sex as a biological variable will require a global shift in science culture. Nature Neuroscience 2021 24:*4*. 2021;24(4):457-464. doi:10.1038/s41593-021-00806-8

62. Gruene T, Flick K, Stefano A, Shea S, Shansky R. Sexually divergent expression of active and passive conditioned fear responses in rats. Elife. 2015;4. doi:10.7554/ELIFE.11352

63. Tan S, Xue S, Behnood-Rod A, et al. Sex differences in the reward deficit and somatic signs associated with precipitated nicotine withdrawal in rats. Neuropharmacology. 2019;160:107756. doi:10.1016/J.NEUROPHARM.2019.107756

64. Kokane SS, Perrotti LI. Sex Differences and the Role of Estradiol in Mesolimbic Reward Circuits and Vulnerability to Cocaine and Opiate Addiction. Front Behav Neurosci. 2020;14:74. doi:10.3389/FNBEH.2020.00074/BIBTEX

65. Wiese B, Wilson-Poe AR. Emerging Evidence for Cannabis’ Role in Opioid Use Disorder. Cannabis Cannabinoid Res. 2018;3(1):179. doi:10.1089/CAN.2018.0022

66. Sagheddu C, Muntoni AL, Pistis M, Melis M. Endocannabinoid Signaling in Motivation, Reward, and Addiction: Influences on Mesocorticolimbic Dopamine Function. Int Rev Neurobiol. 2015;125:257–302. doi:10.1016/BS.IRN.2015.10.004

67. Yamaguchi T, Hagiwara Y, Tanaka H, et al. Endogenous cannabinoid, 2-arachidonoylglycerol, attenuates naloxone-precipitated withdrawal signs in morphine-dependent mice. Brain Res. 2001;909(1-2):121–126. doi:10.1016/S0006-8993(01)02655-5

68. Lichtman AH, Fisher J, Martin BR. Precipitated cannabinoid withdrawal is reversed by Δ9-tetrahydrocannabinol or clonidine. Pharmacol Biochem Behav. 2001;69(1-2):181–188. doi:10.1016/S0091-3057(01)00514-7

69. McPartland JM, Duncan M, Di Marzo V, Pertwee RG. Are cannabidiol and Δ(9) - tetrahydrocannabivarin negative modulators of the endocannabinoid system? A systematic review. Br J Pharmacol. 2015;172(3):737–753. doi:10.1111/BPH.12944

70. Parker LA, Burton P, Sorge RE, Yakiwchuk C, Mechoulam R. Effect of low doses of delta9-tetrahydrocannabinol and cannabidiol on the extinction of cocaine-induced and amphetamine-induced conditioned place preference learning in rats. Psychopharmacology (Berl). 2004;175(3):360–366. doi:10.1007/S00213-004-1825-7

71. Viudez-Martínez A, García-Gutiérrez MS, Medrano-Relinque J, Navarrón CM, Navarrete F, Manzanares J. Cannabidiol does not display drug abuse potential in mice behavior. Acta Pharmacol Sin. 2019;40(3):358. doi:10.1038/S41401-018-0032-8

72. Taylor L, Crockett J, Tayo B, Checketts D, Sommerville K. Abrupt withdrawal of cannabidiol (CBD): A randomized trial. Epilepsy Behav. 2020;104(Pt A). doi:10.1016/J.YEBEH.2020.106938

73. Galaj E, Xi ZX. Possible Receptor Mechanisms Underlying Cannabidiol Effects on Addictive-like Behaviors in Experimental Animals. Int J Mol Sci. 2021;22(1):1–14. doi:10.3390/IJMS22010134

74. de Carvalho CR, Takahashi RN. Cannabidiol disrupts the reconsolidation of contextual drug-associated memories in Wistar rats. Addiction Biology. Published online 2017. doi:10.1111/adb.12366

75. Peres FF, Levin R, Suiama MA, et al. Cannabidiol prevents motor and cognitive impairments induced by reserpine in rats. Front Pharmacol. 2016;7(SEP):343. doi:10.3389/FPHAR.2016.00343/BIBTEX

76. Nedelescu H, Wagner GE, De Ness GL, et al. Cannabidiol Produces Distinct U-Shaped Dose-Response Effects on Cocaine-Induced Conditioned Place Preference and Associated Recruitment of Prelimbic Neurons in Male Rats. Biological Psychiatry Global Open Science. Published online July 7, 2021. doi:10.1016/J.BPSGOS.2021.06.014

77. Linares IM, Zuardi AW, Pereira LC, et al. Cannabidiol presents an inverted U-shaped dose-response curve in a simulated public speaking test. Braz J Psychiatry. 2019;41(1):9–14. doi:10.1590/1516-4446-2017-0015

78. Smith LC, Tieu L, Suhandynata RT, et al. Cannabidiol reduces withdrawal symptoms in nicotine-dependent rats. Psychopharmacology (Berl). 2021;238(8):2201–2211. doi:10.1007/S00213-021-05845-4/FIGURES/5

79. Ożarowski M, Karpiński TM, Zielińska A, Souto EB, Wielgus K. Cannabidiol in Neurological and Neoplastic Diseases: Latest Developments on the Molecular Mechanism of Action. International Journal of Molecular Sciences 2021*, Vol* 22, *Page* 4294. 2021;22(9):4294. doi:10.3390/IJMS22094294

80. Cicero TJ, Ennis T, Ogden J, Meyer ER. Gender differences in the reinforcing properties of morphine. Pharmacol Biochem Behav. 2000;65(1):91–96. doi:10.1016/S0091-3057(99)00174-4

81. Cicero TJ, Aylward SC, Meyer ER. Gender differences in the intravenous self-administration of mu opiate agonists. Pharmacol Biochem Behav. 2003;74(3):541–549. doi:10.1016/S0091-3057(02)01039-0

82. Ramôa CP, Doyle SE, Naim DW, Lynch WJ. Estradiol as a Mechanism for Sex Differences in the Development of an Addicted Phenotype following Extended Access Cocaine Self-Administration. Neuropsychopharmacology 2013 38:*9*. 2013;38(9):1698-1705. doi:10.1038/npp.2013.68

83. Mirbaha H, Tabaeizadeh M, Shaterian-Mohammadi H, Tahsili-Fahadan P, Dehpour AR. Estrogen pretreatment modulates morphine-induced conditioned place preference in ovariectomized mice. Pharmacol Biochem Behav. 2009;92(3):399–403. doi:10.1016/J.PBB.2009.01.009

84. Roth ME, Casimir AG, Carroll ME. Influence of estrogen in the acquisition of intravenously self-administered heroin in female rats. Pharmacol Biochem Behav. 2002;72(1-2):313–318. doi:10.1016/S0091-3057(01)00777-8

85. Fattore L, Fratta W. How important are sex differences in cannabinoid action? Br J Pharmacol. 2010;160(3):544. doi:10.1111/J.1476-5381.2010.00776.X

86. Javadi-Paydar M, Nguyen JD, Kerr TM, et al. Effects of Δ9-THC and cannabidiol vapor inhalation in male and female rats. Psychopharmacology (Berl). 2018;235(9):2541. doi:10.1007/S00213-018-4946-0

87. Fattore L, Spano MS, Altea S, Angius F, Fadda P, Fratta W. Cannabinoid self-administration in rats: sex differences and the influence of ovarian function. Br J Pharmacol. 2007;152(5):795. doi:10.1038/SJ.BJP.0707465

